# “An intrinsically disordered intracellular domain of PIEZO2 is required for force-from-filament activation of the channel”

**DOI:** 10.1101/2021.01.13.426495

**Authors:** Clement Verkest, Irina Schaefer, Juri M. Jegelka, Timo A. Nees, Wang Na, Francisco J. Taberner, Stefan G. Lechner

## Abstract

A central question in mechanobiology is how mechanical forces acting in or on a cell are transmitted to mechanically-gated PIEZO channels that convert these forces into biochemical signals. Here we show that PIEZO2 is sensitive to force-transmission via the membrane (force-from-lipids) as well as force transmission via the cytoskeleton (force-from-filament) and demonstrate that the latter requires the intracellular linker between the transmembrane helices nine and ten (IDR5). Moreover, we show that rendering PIEZO2 insensitive to force-from-filament by deleting IDR5 abolishes PIEZO2-mediated inhibition of neurite outgrowth, which relies on the detection of cell-generated traction forces, while it only partially affects its sensitivity to cell indentation and does not at all alter its sensitivity to membrane stretch. Hence, we propose that PIEZO2 is a polymodal mechanosensor that detects different types of mechanical stimuli via different force transmission pathways, which highlights the importance of utilizing multiple complementary assays when investigating PIEZO channel function.

## Main text

Virtually all cells of our organism are constantly exposed to mechanical forces of one kind or another. Thus, besides the ubiquitous gravitational and osmotic forces, some cells experience compression and stretch induced by externally applied mechanical stimuli or by organ distension, respectively, while others are exposed to shear stress exerted by circulating body fluids. Moreover, many cells generate traction forces acting on their surface when they explore their local environment. Accordingly, most cells are equipped with sensors that enable them to detect and convert mechanical stimuli into biochemical signals – a process called mechanotransduction – that trigger adaptive processes required to maintain cell, tissue and not least body integrity in an everchanging mechanical environment.

Since their discovery in 2010^1^, the mechanically-activated ion channels PIEZO1 and PIEZO2 were shown to be of crucial importance for mechanotransduction in a variety of tissues. PIEZO1, for example, detects shear stress and stretch in erythrocytes, vascular endothelial cells, chondrocytes and bladder urothelial cells^2,3^. PIEZO2, on the other hand, appears to be particularly important for mechanotransduction in primary sensory afferents as it was shown to be involved in the detection of light touch, mechanical pain, proprioception, airway stretch and bladder distension^4–11^. Moreover, PIEZOs contribute to the regulation of processes such as neurite outgrowth^12^, wound healing^13^ and tumor cell dissemination^14^, probably by detecting cell-generated traction forces acting on the plasma membrane during neurite extension and cell migration^15–17^.

Recent high-resolution cryo-EM studies showed that PIEZO1 and PIEZO2 oligomerize as homotrimers with a propeller-shaped quaternary structure that locally curve the membrane into a spherical dome^18–21^. The “propeller” blades are formed by 36 transmembrane helices that are grouped into nine four-transmembrane-helixcontaining bundles termed ‘Piezo repeats’ or transmembrane helical units (THUs) and are thought to be involved in sensing membrane tension^22–24^. The ion conducting pore is formed by three transmembrane helices (one from each protamer) termed inner helices and the intracellular C-terminal domains (CTD). In addition to the central pore, PIEZOs also comprise three lateral ion-conducting portals that branch off the main ion conduction pathway and that are gated by the spliceable plug domains^25,26^. Two other peculiar domains that are important for PIEZO channel function are the so-called ‘cap’ domain and the ‘beam’ domain. The cap, which is also called C-terminal extracellular domain (CED), is located on the extracellular side above the central ion conducting pore and is coupled to the blades by electrostatic interactions, which together with hydrophobic amino acids in the pore lining inner helices determine the inactivation kinetics of PIEZO channels^27–29^. The beam domain is a long α-helix that protrudes from the intracellular side of THU7 toward the inner vestibule of the channel and is thought to contribute to channel activation by relaying conformational changes of the blades to the pore-forming domains^30^.

However, despite our profound understanding of the roles of different PIEZO domains in channel activation and inactivation, a fundamental question that remains open is how mechanical forces that act on the cell surface are transmitted to the channel in the first place. Two paradigms are commonly used to explain mechanical force-induced channel activation. The force-from-lipids model proposes that mechanically-induced membrane tension causes changes in the transbilayer pressure profile asymmetry, which leads to conformational changes and thus activation of PIEZOs^31–33^. While there is compelling evidence that PIEZO1 is activated by force-from-lipids under experimental conditions where the cytoskeleton is completely missing (e.g. droplet bilayers, excised patches, membrane blebs)^24,34,35^, it is still unclear if this activation mechanism also applies to PIEZO2. Moreover, it is unclear how PIEZOs are activated in intact cells where the lipid bilayer is bound to the underlying actin cortex by various transmembrane proteins and additional mechanism might thus come into play. Indeed, it was recently shown that changes in membrane tension are locally restricted and are not propagated across the cell surface when the membrane is bound to the cytoskeleton^36^, which raises the question as to if and how channels that are located further away from the site of mechanical stimulation are activated. Such “long-distance” activation of PIEZOs can be explained by the so-called force-from-filament model, which proposes that PIEZOs are tethered to the cytoskeleton, such that mechanically-induced movements of the cytoskeleton activate the channel by pulling or pushing it open from the intracellular side. While intracellular tethering was convincingly demonstrated for other mechanosensitive channels (e.g. NOMPC, TMC1^37,38^), there is only indirect evidence supporting force-from-filament gating of PIEZO channels. Thus, several studies showed that pharmacological disruption of the cytoskeleton with Cytochalasin-D dramatically reduces membrane indentation-evoked whole-cell PIEZO currents^39,40^. While these studies clearly showed that an intact cytoskeleton is essential for efficient PIEZO activation, they did not clarify whether cytoskeletal tension is directly transferred to PIEZOs via tethers (i.e. force-from-filament) or whether it solely causes changes in local membrane tension that ultimately activate the channel via force-from-lipids^31,32^.

Interestingly PIEZO channels comprise several large intracellular domains that are structurally unresolved and the function of which is hitherto completely unclear. Considering the remarkable size of these intrinsically disordered regions (IDRs), which in PIEZO2 account for ~25% of the channel, together with the fact that they are ideally located to mediate interactions between PIEZO2 and the cytoskeleton, which might be required for force-from-filament gating, we here set out to examine their role in PIEZO2 function.

## Results

### Deletion of IDR5 reduces membrane indentation-evoked but not membrane stretch-evoked PIEZO2 currents

PIEZO2 comprises seven large intracellular and intrinsically disordered regions (IDRs) that together account for ~25% of the amino acids sequence of the channel (Fig. 1a and b). IDR1, 4, 5, 6 and 7 serve as linkers between adjacent THUs, whereas IDR2 and IDR3 link THU8 with the clasp domain and the clasp with the latch domain, respectively (Supplementary Fig. 1a). To examine the role of these IDRs we generated PIEZO2 mutants that lack individual IDRs (IDR1^del^ – IDR7^del^, Fig. 1b and Supplementary Fig. 1a) and assessed their functional properties in Neuro2a-PIEZO1-KO (N2a) cells, which completely lack endogenous mechanotransduction currents^24^. We first compared PIEZO2 and IDR1^del^-IDR7^del^-mediated transmembrane currents evoked by mechanical indentation of the cell membrane using the whole-cell configuration of the patch-clamp technique (Fig. 1c), which is the most commonly used experimental approach for studying PIEZO2 function. Strikingly, only deletion of IDR3 and IDR5 altered PIEZO2 function in this assay. Thus, IDR5^del^-mediated currents exhibited dramatically reduced amplitudes (IDR5: 0.34 ± 0.07 nA vs. PIEZO2 1.13 ± 0.12 nA at 5.2 μm indentation, Fig. 1c and d) and slightly, yet significantly, increased activation thresholds (Fig. 1e). The inactivation kinetics and the reversal potential were, however, not affected by the deletion of IDR5 (Fig. 1f and g, Supplementary Fig. 2). By contrast, IDR3^del^-mediated currents were more than twice as big as full-length PIEZO2-mediated currents (IDR3^del^: 2.6 ± 0.43 nA vs. PIEZO2 1.13 ± 0.12 nA at 5.2 μm Fig. 1c and d), while exhibiting similar activation thresholds, inactivation kinetics and reversal potentials (Fig. 1e-g, Supplementary Fig. 2). All other IDR-deletions produced mechanical indentation-evoked currents that were indistinguishable from those mediated by full-length PIEZO2 (Fig. 1d-g, Supplementary Fig. 2).

**Fig. 1.**
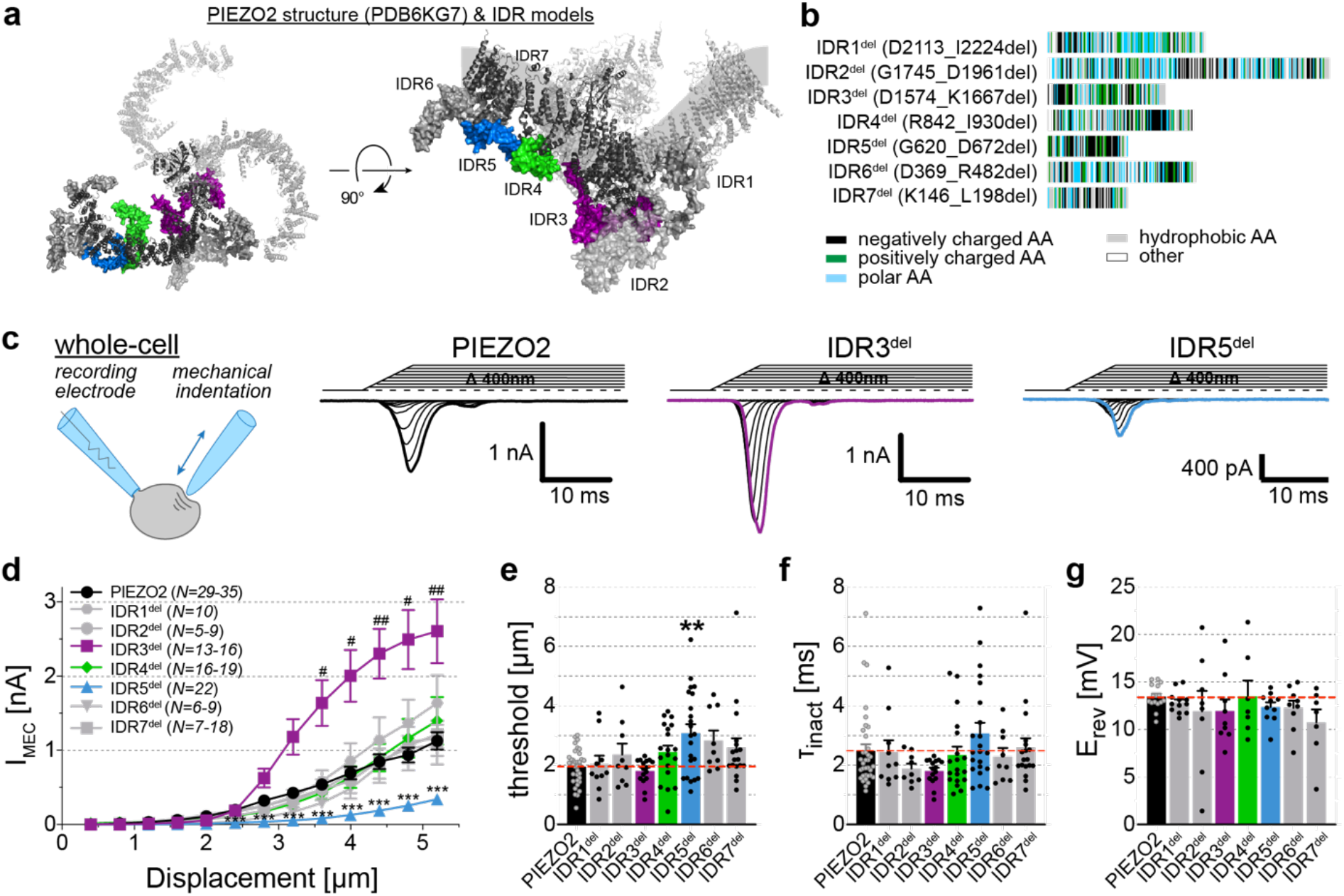
Deletion of IDR3 and IDR5 alters membrane indentation-evoked PIEZO2 currents. **a**, Top (left) and side (right) view of the mouse PIEZO2 structure (PDB6KG7). Structural models of the IDR regions (1-7, in gray, blue, green or purple) were added along one of the three PIEZO2 blades (black). **b**, Amino acid position and color-coded properties of the IDRs that have been deleted. **c**, Cartoon depicting the recording paradigm (left) and representative example traces (right) of whole-cell mechanically activated currents evoked by increased probe displacement of PIEZO2 and indicated IDR^del^ mutants, recorded at −60 mV. **d**, Displacement-responses curves of peak current amplitudes of PIEZO2 and IDR^del^ mutants in N2a-P1KO cells. Symbols represent means ± s.e.m. Comparison with Kruskal-Wallis test and Dunn’s post-test (* PIEZO2 vs IDR5^del^; # PIEZO2 vs IDR3^del^). *n* numbers of cells per group are indicated in the graph legend. **e**, Comparison of the mechanical activation thresholds of PIEZO2 and IDR^del^ mutants performed with One-Way ANOVA, F(7,130)=4.093, Dunnett’s post-test P2WT vs IDR5^del^, ***p<0.0001. **f**, Comparison of the inactivation time constants (τ_inact_) of PIEZO2- and IDR^del^-mediated currents using the Kruskal-Wallis test (ns p=0.1531). For **e-f**, bars represent means ± s.e.m., with individual values. *N-number* per group are identical to those in **c**. **g**, Reversal potential (E_rev_) of PIEZO2 and IDR^del^-mutants. Bars represent means ± s.e.m., with individual values. Comparison with Kruskal-Wallis, ns p=0.4886. *n* per group are (from left to right): 16, 12, 8, 9, 7, 10, 9, 7.

Another experimental approach for studying PIEZO channel function is the so-called pressure-clamp technique. Here, the currents are recorded in the cell-attached mode of the patch-clamp technique and the channels are activated by stretching the membrane patch inside the patch-pipette by application of negative pressure (Fig. 2a). Unlike PIEZO1, which is robustly activated by pressure-induced membrane stretch in cell-attached recordings^1,24,34^, PIEZO2 only rarely responds to such stimuli^18,24,41,42^. Consistent with previous studies that reported pressure-induced currents in ~20% of PIEZO2 expressing cells, we observed stretch-evoked PIEZO2 currents in 8 from 38 recorded cells (Fig. 2a and b). Interestingly, the proportion of cells that exhibited pressure-induced currents was more than twice as big amongst cells expressing IDR1^del^ (9/19 cells), IDR2^del^ (8/18 cells), IDR4^del^ (25/42 cells) and IDR5^del^ (10/22 cells), but this difference was only statistically significant for IDR4^del^ (Fig. 2b). Since the pressure-evoked PIEZO2 currents never saturated in the pressure range in which the recordings were stable (note that pressures below – 80 mmHg frequently ruptured the membrane) and, moreover, did not exhibit clearly discriminable single channel openings or peaks at higher pressures (Fig. 2a), we were unable to estimate the total number of channels in the patch and thus could not calculate dwell times and open probabilities. Hence, in order to statistically compare the pressure-evoked currents, we determined the total charge transfer by measuring the area under the curve (AUC) over the time of the pressure stimulus. This analysis revealed, that IDR4^del^ did not only respond more frequently to pressure-induced membrane stretch, but also generated significantly larger currents (PIEZO2: 1.11 ± 0.41 pC vs. IDR4: 4.05 ± 1 pC at −60 mmHg, Fig. 2c).

**Fig. 2.**
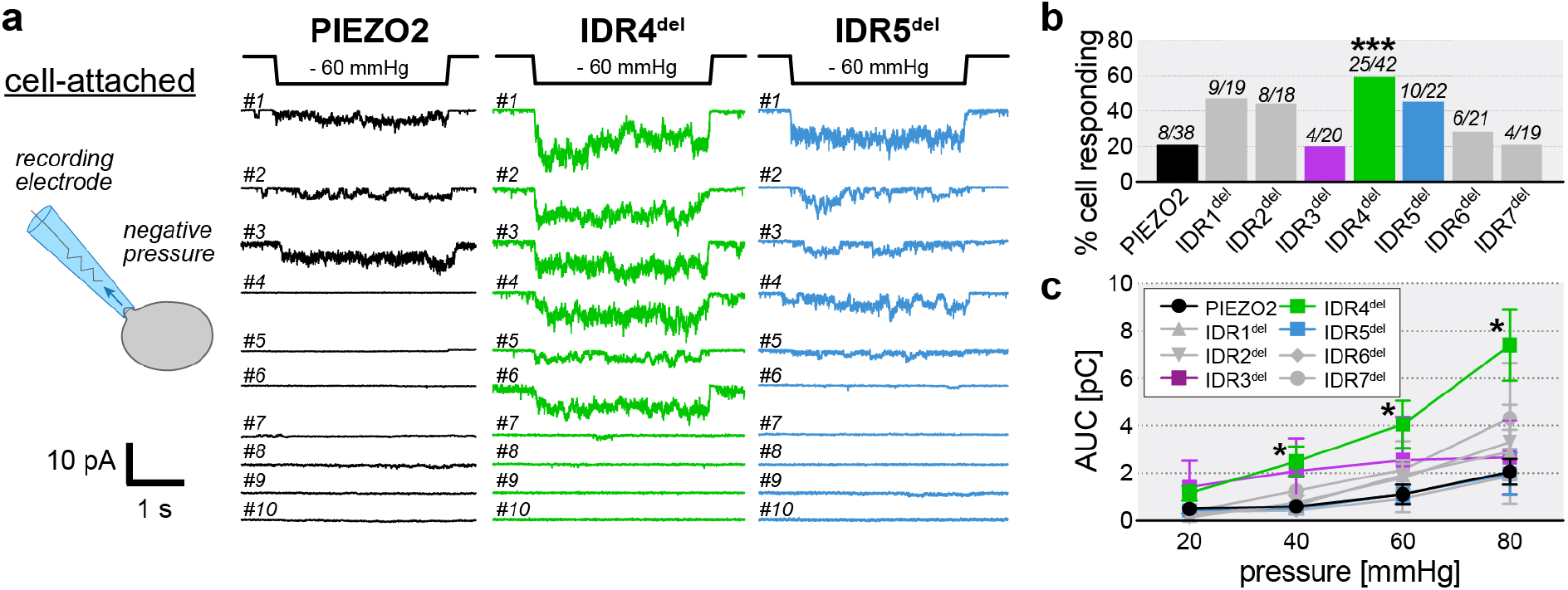
Deletion of IDR4 increases stretch-sensitivity of PIEZO2. **a**, Cartoon depicting the recording paradigm (left) and example traces of the indicated stretch-activated cell-attached currents at −100 mV evoked by a −60 mmHg negative pressure pulse (3 s duration) of PIEZO2 and the indicated IDR^del^ mutants from 10 different cells. **b**, Comparison of the proportion of cells responding to stretch in PIEZO2 and individual IDR^del^ mutant. Fisher’s exact test, PIEZO2 vs IDR1^del^ ns p=0.0646, PIEZO2 vs IDR4^del^ *** p=0.0006, PIEZO2 vs IDR5^del^ ns p=0.07801. The total number of cells recorded and the number of responders are indicated above each bar. **c**, Pressure-response curves of total charge transferred (AUC) by PIEZO2 and IDR^del^ mutants in N2a-P1KO cells. Symbols represent means ± s.e.m. Comparison with Kruskal-Wallis test (p=0.0569 −20 mmHg, p=0.0222 −40 mmHg, p=0.0037 −60 mmHg, p=0.0027 −80 mmHg) and Dunn’s post-test, PIEZO2 vs IDR4^del^ (* p=0.0427 −40 mmHg, * p=0.0427 −60 mmHg, * p=0.0483 −80 mmHg). *n* number of cells per group is indicated in the graph legend.

To test if changes in the unitary conductance underlie the large stretch-evoked IDR4^del^-currents and the altered poking-evoked IDR3^del^- and IDR5^del^-current amplitudes, we next measured the single channel current amplitudes at different holding potentials and calculated the unitary conductance by linear regression of the current-voltage relationships (Fig. 3a-d). This analysis showed that IDR3^del^ has a significantly larger single channel conductance than full-length PIEZO2 (IDR3^del^, 34.07 ± 1.26 pS vs.

**Fig. 3.**
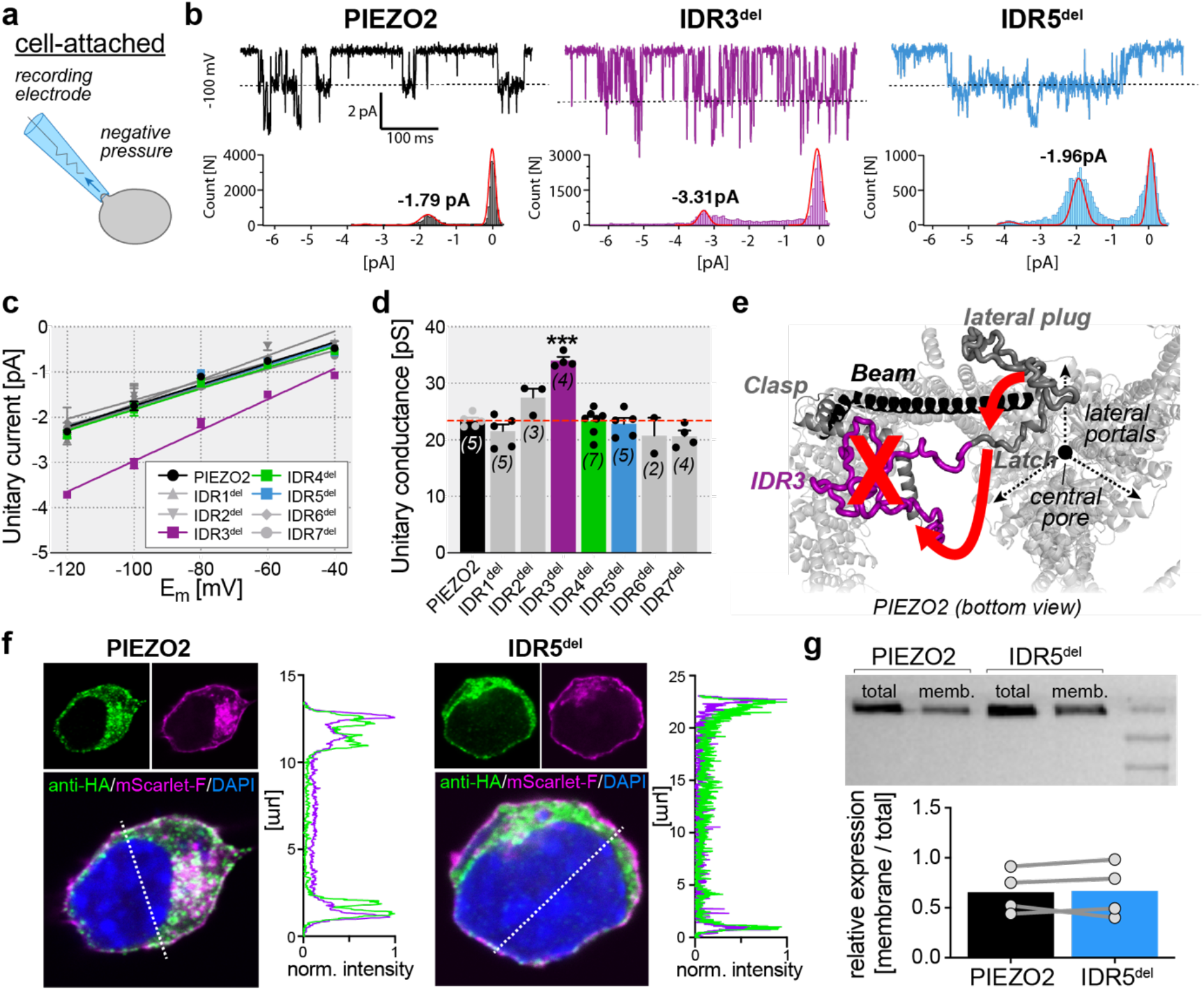
Ion permeation and membrane expression of PIEZO2 and IDR^del^ mutants. **a**, Cartoon depicting the recording paradigm. **b**, Representative example traces of stretch-activated single-channel currents at −100 mV evoked by negative pressure stimuli of PIEZO2 and indicated IDR^del^ mutants (top), and corresponding histogram analysis of the single channel recorded segment (bottom). The value of the peak of the Gaussian fit (in pA) is indicated. **c**, Linear regression of the I-V relationships of the indicated PIEZO2 constructs. Symbols are means ± s.e.m. *n* number are the same is in **d**. **d**, Unitary conductance of PIEZO2 and IDR^del^ mutants. Bars represent means ± s.e.m. Comparison with Kruskal-Wallis (p=0.0110) and Dunn’s post-test, P2WT vs IDR3^del^, * p=0.0104. *n*-numbers are indicated in the graph. **e**, Proposed model explaining the increased currents of IDR3^del^. Bottom view of PIEZO2 structure, focused on the pore axis. Removal of IDR3 (purple) might pull away (red arrow) the lateral plug and the latch (dark grey) and stretch them to connect back (red arrow) to the Clasp located at the other side of the Beam (black), ultimately resulting in the opening of the lateral portals. **f**, Representative immunofluorescence images of PIEZO2 (left) and IDR5^del^ (right) subcellular localization. Transfected N2A-P1KO were stained for PIEZO (HA tag, green), nucleus (DAPI, blue) and plasma membrane was visualized with endogenous mScarlet (magenta). Graphic adjacent is the intensity of the normalized fluorescence along the dotted white line in the PIEZO (HA) and membrane (mScarlet) channel. **g**, Western blots (top) of biotinylated membrane fraction and whole-cell lysate samples (total fraction) derived from N2A-P1KO cells transfected with PIEZO2 or IDR5^del^ and densitometric quantification (bottom) of the membrane fraction compared to the total one in PIEZO2 and IDR5^del^. Bars show the means with individual values from 4 independent assays. Connecting lines represent the paired experimental values. Mann-Whitney test, ns p=0.7553.

PIEZO2, 23.4 ± 1.14 pS, mean ± SEM), whereas all other IDR-deletions, including IDR4^del^ and IDR5^del^, have single channel conductances that are in the same range (Fig. 3b-d). IDR3 links the N-terminal side of the clasp domain to the C-terminal end of the latch domain (Supplementary Fig. 1), which on its N-terminal side is connected to the beam domain by a 44 amino acid long linker (Fig. 3e). We had previously shown that the beam-to-latch linker controls ion permeation of PIEZO2 and PIEZO1^26^ and Geng et al. later found that it does so by acting as a plug that blocks the lateral portals of PIEZO channels, which – when unblocked – contribute to ion permeation in addition to the central pore^25^. Considering that IDR3 spans a distance of approximately 7 nm, we hypothesize that deletion of IDR3 causes dislocation of the adjacent lateral plug, such that the lateral portals are unblocked and single channel conductance is increased (Fig. 3e).

Since the unitary conductance of PIEZO2 was not affected by the deletion of IDR5 (Fig. 3d), we next considered the possibility that altered membrane trafficking could account for the reduced poking-evoked current amplitudes of IDR5^del^. To address this question, we co-transfected N2a cells with a membrane bound farnesylated variant of the fluorescent protein mScarlet and full-length PIEZO2 and IDR5^del^, respectively, and visualized the subcellular localization of the proteins with immunocytochemistry. As the line scans of the confocal sections in Figure 3f show, the fluorescence signals of both full-length PIEZO2 as well as IDR5^del^ overlap with the membrane bound mScarlet fluorescence, indicating that IDR5^del^ is properly trafficked to the plasma membrane. To corroborate this observation, we also quantified cell-surface expression of PIEZO2 and IDR5^del^ using biotinylation and subsequent western blot analysis. Again, we found no differences in the membrane expression levels of IDR5^del^ compared to full-length PIEZO2 (Fig. 3g). These results together with the observation that almost 50% of the IDR5^del^-expressing cells exhibited pressure-evoked currents (Fig. 2a and b) – which clearly demonstrates the presence of functional channels in the plasma membrane – strongly suggest that altered membrane trafficking does not account for the small poking-evoked current amplitudes of IDR5^del^.

In summary, the characterization of the basal properties of the IDR-deletions provided two notable insights. Firstly, the finding that deletion of IDR3 increases the single channel conductance of PIEZO2, which we believe is caused by dislocation of the adjacent beam-to-latch linker, supports previous studies that have suggested that this domain forms a plug that prevents ion permeation through the lateral portals of PIEZO channels. Secondly, the observation that deletion of IDR4 increases stretch-sensitivity but does not affect poking-sensitivity, while the deletion of IDR5 has the opposite effect – i.e. reduced poking-sensitivity but normal stretch-sensitivity – suggests that poking and stretch activate PIEZO2 via two different mechanisms.

### Deletion of IDR5 prevents PKA-dependent modulation of PIEZO2

PIEZO2 has been shown to be modulated by several G-protein-coupled receptors (GPCRs) such as the P2Y2-receptor, the Bradykinin B2 receptor and the GABAB receptor, as well as by direct activation of the classical GPCR downstream effectors Protein Kinase A and C (PKA, PKC)^43–45^. Since most IDRs contain numerous consensus PKA phosphorylation sites (Fig. 4a and Supplementary Fig. 1a), we hypothesized that in addition to controlling the basal properties of PIEZO2, some IDRs might also be required for the PKA-dependent modulation of PIEZO2. To test this hypothesis, we compared mechanical indentation-evoked currents recorded in the presence of the PKA inhibitor KT5720 with currents recorded from cells treated with the PKA activator 8-Br-cAMP. In addition, all cells were treated with the PKC inhibitor GF109203X to avoid a possible confounding influence from basal PKC activity. As shown in Figure 4b and c, PKA activation significantly increased the amplitudes of fulllength PIEZO2-mediated currents, which was consistent with a previous report describing the PKA-dependent modulation of PIEZO2^45^. Strikingly, all IDR-deletions except for IDR5^del^ were modulated to the same extent as full-length PIEZO2 – i.e. significantly increased current amplitudes. Consistent with the increase in current amplitudes, there was also a trend towards lower thresholds (Fig. 4d) and, interestingly, shorter inactivation time constants (Fig. 4e), in all IDR-deletions except for IDR5. Considering that the only IDR that appeared to be required for PKA-dependent modulation of PIEZO2 – i.e. IDR5 – does not contain a single consensus PKA site (Fig. 4a and Supplementary Fig. 1a), our data suggest that the PKA-dependent potentiation of PIEZO2 observed in our expression conditions does not require phosphorylation of the channel and is probably mediated by an indirect mechanism.

**Fig. 4.**
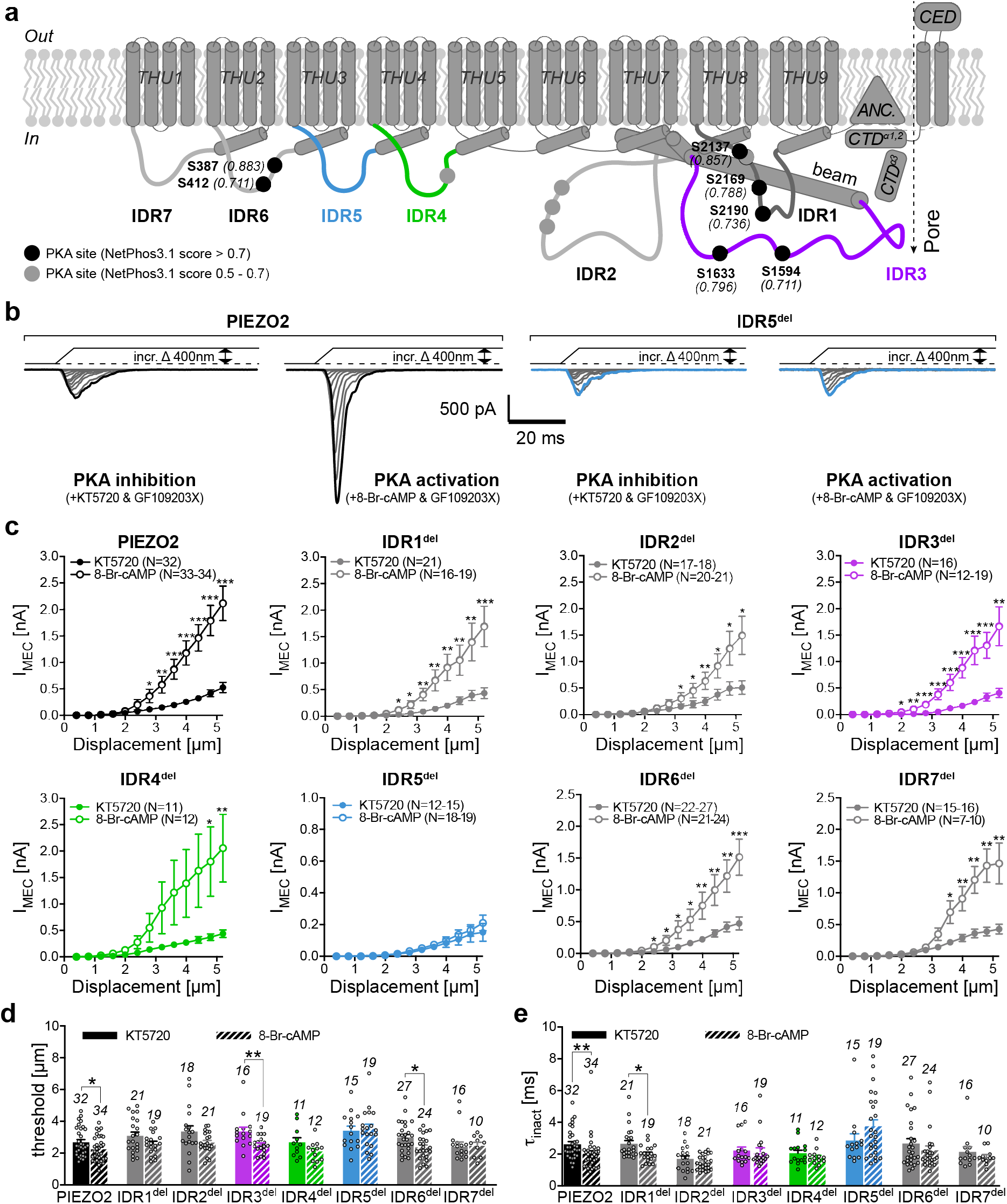
IDR5 is required for PKA-dependent modulation of PIEZO2. **a**, Membrane topology of PIEZO2 with its major domains and the predicted PKA phosphorylation sites (grey and black circles). **b**, Example traces of mechanically activated currents evoked by increased probe displacement of PIEZO2 (left, black) and IDR5^del^ (right, blue) after PKA/PKC inhibition (KT5720 and GF109203X, 1μM each) and PKA activation (300μM 8-Br-cAMP and 1 μM GF109203X). **c**, Comparison of the displacement-responses curves of peak current amplitudes of PIEZO2 and IDR^del^ mutants after inhibition and activation of PKA. All data points represent means ± s.e.m. and *N*-numbers of cells per group are indicated in the graph legends. Data points were compared using Mann-Whitney test. **d**, Comparison of the mean ± s.e.m. mechanical activation thresholds of the indicated mutants after inhibition (solid bars) and activation (striped bars) of PKA. Mann-Whitney test, GF+KT vs 8-Br-cAMP+GF: PIEZO2 * p=0.0389, IDR3^del^ ** p=0.0018, IDR6^del^ * p=0.0284. **e,** Inactivation time constant (τ_inact_) of PIEZO2 and IDR^del^ after inhibition (solid bars) and activation of PKA (striped bars). Mann-Whitney test, GF+KT vs 8-Br-cAMP+GF: PIEZO2 ** p=0.001, IDR1^del^ *p=0.0126.

### IDR5 is required for force-from-filament gating of PIEZO2

PKA is known to play a crucial role in the remodeling of the cytoskeleton^46^, which was shown to be involved in the activation of membrane indentation-evoked PIEZO currents – presumably by transmitting mechanical force from the site of stimulation to channels that are located further away^31,32^. Considering that IDR5^del^ was not efficiently activated by membrane indentation (Fig. 1c and d) and considering further that it was not potentiated by PKA (Fig. 4b and c), we hypothesized that IDR5 might serve as an interface that mediates force transmission from the cytoskeleton to PIEZO2.

To test this hypothesis, we examined the effect of the actin cytoskeleton-disrupting drug Cytochalasin-D on PIEZO2 and IDR5^del^-mediated currents. As previously shown for PIEZO1^35,40,47^ and PIEZO2^39,48^, treatment with Cytochalasin-D (15 min, 1 μM) significantly reduced membrane indentation-evoked current amplitudes of full-length PIEZO2 (Fig. 5a and b) without affecting the mechanical activation threshold and inactivation kinetics (Fig. 5c and d). By contrast, pressure-induced PIEZO2 currents in cell-attached recordings, tended to be larger and were more frequent after Cytochalasin-D treatment (Fig. 5 e-g), which was consistent with previous reports describing similar effects of Cytochalasin-D treatment on PIEZO1 currents^35,40^. Inhibition of microtubule polymerization with Nocodazole (30 min, 1 μM) neither affected stretch-nor poking-evoked PIEZO2 currents (Fig. 5a-g). Most importantly, disruption of the actin cytoskeleton did not reduce membrane indentation-evoked IDR5^del^ currents (Fig. 5h-k), demonstrating that the activation of IDR5^del^ does not require an intact cytoskeleton.

**Fig. 5.**
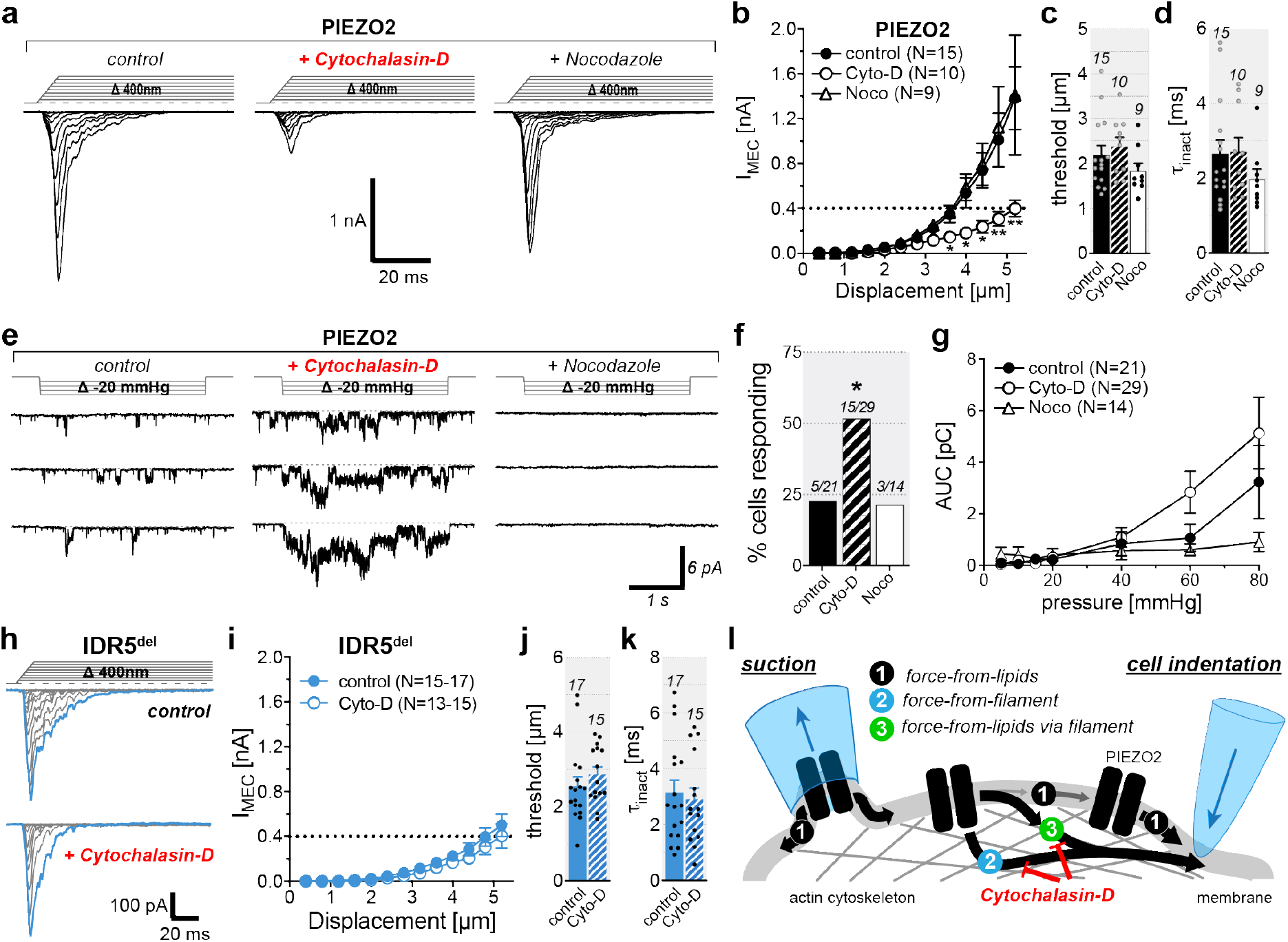
Actin cytoskeleton disruption impairs PIEZO2-but not IDR5^del^-mediated membrane indentation-evoked currents. **a,** Example traces of mechanically activated PIEZO2 currents evoked by increased probe displacement under control conditions (left), after treatment with Cytochalasin-D (middle) and after Nocodazole treatment (right). **b**, Displacement-responses curves of peak current amplitudes of PIEZO2 in the indicated conditions. Symbols represent means ± s.e.m. and date were compared using Kruskal-Wallis test followed by Dunn’s post-test (* Control vs Cytochalasin-D). *n*-numbers of cells per group are indicated in graph legend. **c**, Comparison of the mechanical activation thresholds of PIEZO2 in the indicated conditions using Kruskal-Wallis test (p=0.1725). **d,** Comparsion of the inactivation time constants (τ_inact_) of PIEZO2-currents recorded in the indicated conditions using Kruskal-Wallis test (p=0.3148). Bars in **c-d** represent means ± s.e.m., with individual values shown as circles. N-numbers per group are indicated above each bar**. e,** Representative example traces of PIEZO2 stretch-activated currents recorded under the conditions indicated above the traces at −100 mV and evoked by increasing negative pressure pulses. **f**, Comparison of the proportinos of cells responding to stretch. Fisher’s exact test, PIEZO2-control vs PIEZO2-Cytochalasin-D * p=0.0441. *n*-numbers of cells are indicated above the bars. **g**, Pressure-responses curves of PIEZO2-control and with Cytochalasin-D or Nocodazole. Symbols are means ± s.e.m. Comparison with Kruskal-Wallis test. *N*-number of cells per group are indicated in the graph legend. **h,** Example traces of mechanically activated currents evoked by increased probe displacement of IDR5^del^ (top) and in the presence of Cytochalasin-D (bottom). **i**, Displacement-responses curves of IDR5^del^ in the presence or absence of Cytochalasin-D. Symbols are mean ± s.e.m. Comparison with Mann-Whitney test. *n* number of cells per group is indicated in graph legend. **j**, Displacement threshold of mechanically activated currents mediated by IDR5^del^ without (blue) or with Cytochalasin-D (stripped). Mann-Whitney test, p=0.2304. **k,** Inactivation time constant (τ_inact_) of IDR5^del^ with or without Cytochalasin-D. Unpaired-t test, p=0.6933. Bar graphs in **j-k,** are mean ± s.e.m., with individual values. *n* per group are above each bar. **l,** Cartoon depicting the possible gating mechanisms of PIEZO2 upon suction (left) or cell indentation (right).

For PIEZO1, the diametrically opposing effects of Cytochalasin-D treatment on stretch- and poking-evoked currents are usually explained by the following model^31,32,35^. Since the cytoskeleton protects the plasma membrane from being excessively stretched in response to pressure application – an effect known as mechanoprotection – it is thought that disruption of the cytoskeleton renders the membrane more sensitive to pressure, such that a given pressure pulse results in greater membrane tension and thus larger currents in cell attached recordings. The fact that PIEZO1 can be activated in the complete absence of the cytoskeleton^24,34,35^ together with the observation that disruption of the cytoskeleton does not prevent stretch-activation of the channel^35,40^, has led to the currently prevailing view that PIEZOs are directly activated by force-from-lipids in cell-attached patch-clamp recordings (Fig 5l). The inhibitory effect of cytoskeleton disruption on membrane indentation-evoked currents, on the other hand, is explained by a loss of force transmission from the site of mechanical indentation to channels that are located further away, which is thought to be mediated by the cytoskeleton because locally induced changes in membrane tension are not propagated across the cell surface via the plasma membrane^36^. What is still unclear, however, is how the cytoskeleton ultimately transmits forces to PIEZO channels. While some researchers proposed that cytoskeletal strain causes local membrane tension around the channels such that the channels are ultimately activated by force-from-lipids (i.e. force-from-lipids via filament in Fig. 5l), others believe that PIEZOs are tethered to the cytoskeleton such that cytoskeletal strain directly pulls the channel open – i.e. force-from-filament (Fig. 5l)^31–33^. The former hypothesis implies that a channel that is normally sensitive to membrane stretch, should also be normally sensitive to membrane indentation in cells with an intact cytoskeleton and that poking-evoked currents of such a channel should be inhibited by disruption of the cytoskeleton. IDR5^del^, which does exhibit normal stretch sensitivity, however, fulfils neither of these conditions. Hence, we propose that PIEZO2 is directly activated by force-from-filament and suggest that IDR5 mediates the interaction between the cytoskeleton and the channel.

### PIEZO2 mediated inhibition of neurite outgrowth requires force-from-filament gating

The identification of a PIEZO2 mutant – i.e. IDR5^del^ – that exhibits normal force-from-lipids sensitivity but is insensitive to force-from-filament, enabled us to examine which of the two gating mechanisms is relevant for the detection of naturally occurring stimuli in fully intact cells. An important function of PIEZO channels that has only recently come to light is the detection of cell-generated traction forces that occur, for example, when cells explore their environment or during cell migration and neurite outgrowth^12,14–17^. The N2a cells that we have used throughout this study are a frequently used model system for studying neuronal differentiation and neurite outgrowth. We thus next asked if PIEZO2 has the ability to modulate neurite outgrowth from these cells and if so, how the cell-generated forces activate the channel.

Consistent with previous reports^49,50^, we found that 24 hours treatment with nerve growth factor (NGF, 100 ng/ml) and serum deprivation significantly increased the number of neurite-bearing cells from 41% (343/836 cells) to almost 87% (741/854 cells) amongst cells transfected with a plasmid encoding green fluorescent protein (GFP, Fig. 6a-c). Notably, NGF-treatment more than doubled the mean neurite length from 13.6 ± 0.5 μm (mean ± SEM, N=343) to 28.7 ± 0.8 μm (mean ± SEM, N = 743) and significantly increased the average number of neurites per cell, as illustrated by the cumulative neurite number frequency distribution (Fig. 6a-c). Interestingly, in cells expressing full-length PIEZO2, NGF-treatment and serum-deprivation only had a negligible effect on neurite outgrowth. Thus, although the number of neurite-bearing cells increased to 73 % (733/998 cells) with NGF-treatment, the neurites of PIEZO2-expressing cells were only slightly longer (17.4 ± 0.5 μm, N=733) than those of untreated GFP-expressing cells and significantly shorter than those of NGF-treated and serum deprived GFP-expressing cells. Moreover, the number of neurites per cell was also smaller amongst PIEZO2-expressing cells as compared to GFP-expressing cells (Fig. 6c). Together these results clearly demonstrate that PIEZO2 inhibits neurite outgrowth from N2a cells, though at present we can only speculate about the exact mechanism underlying this inhibition. However, given the complete absence of other mechanical stimuli in this assay, it seems likely that PIEZO2, as previously shown for PIEZO1, generates Ca^2+^ signals in response to cell-generated traction forces that occur during neurite outgrowth. In order to test whether these traction forces activate PIEZO2 via force-from-filament or via force-from-lipids, we next examined neurite outgrowth inhibition by IDR5^del^, which is insensitive to force-from-filament (Fig. 1 and 5) but appears to respond normally to force-from-lipids (Fig. 2 and 3). Strikingly, NGF-treated and serum-deprived IDR5^del^-expressing cells were almost indistinguishable from GFP-expressing cells with respect to neurite length (26.3 ± 0.8 μm, N=685, Fig. 6a and b), number of neurites per cell (Fig. 6a and c) and proportion of neurite-bearing cells (84%, 683/810 cells). Hence, IDR5 appears to be required for the detection of cell-generated traction forces, suggesting that such forces activate PIEZO2 via force-from-filament gating.

**Fig. 6.**
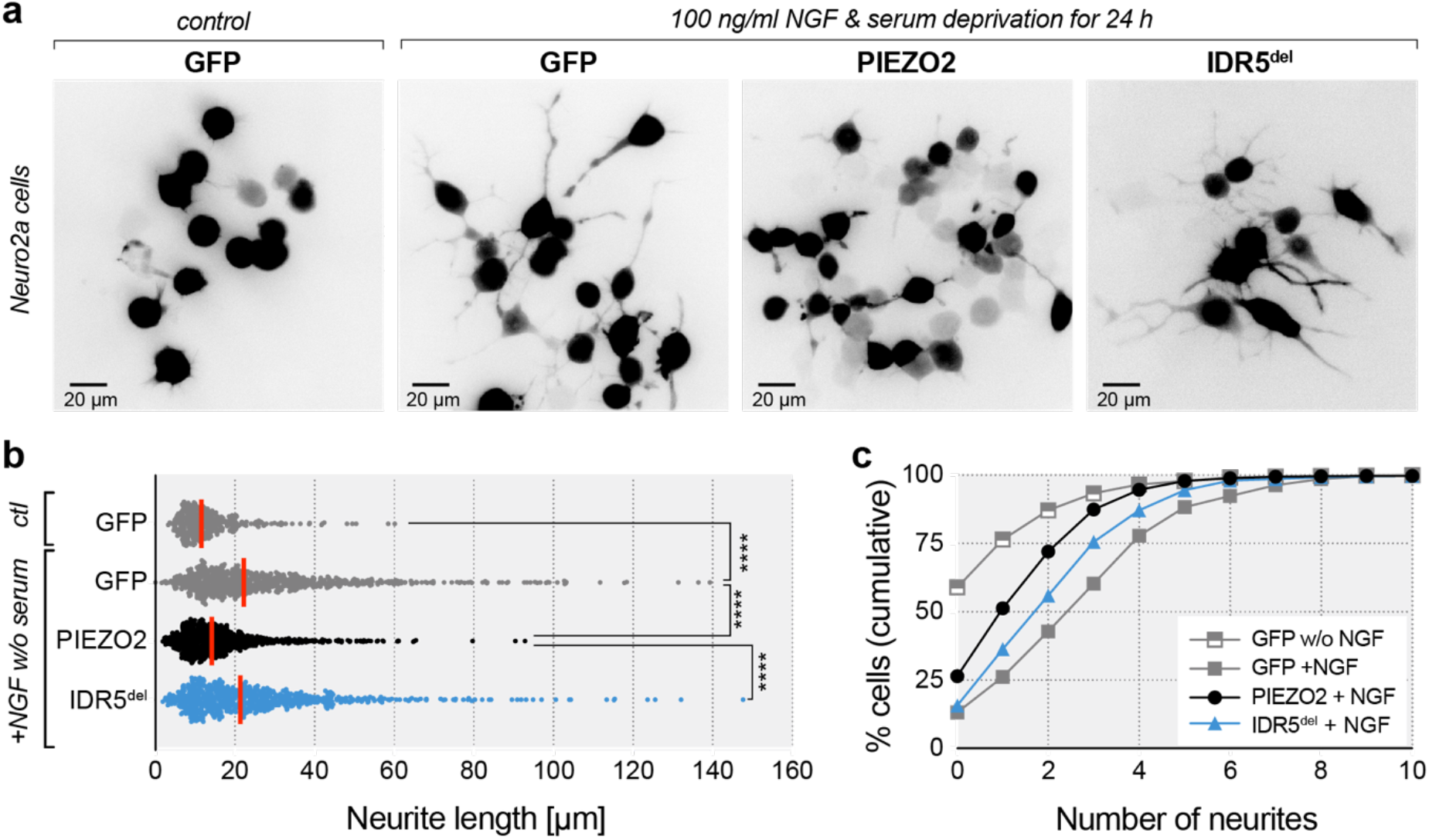
PIEZO2 but not IDR5^del^ inhibits neurite outgrowth from N2a cells. **a,** Representative images showing GFP fluorescence N2a-P1KO cells transfected with GFP only (left and second from left), PIEZO2-HA-IRES-GFP (PIEZO2) and P2-IDR5-IRES-GFP (IDR5^del^), cultured for 24 h in regular growth medium (control) or in the presence of 100 ng/ml NGF and the absence of serum (GFP second from left, PIEZO2 and IDR5^del^). **b**, Scatter dot-plot showing the mean neurite lengths (red lines) as well as the individual values from all cells (scattered symbols) observed under the indicated conditions. Data were compared using the Kruskal-Wallis test, p<0.0001, and Dunn’s post-test: GFP control vs. GFP +NGF w/o serum p<0.0001, GFP +NGF w/o serum vs. PIEZO2 p<0.0001, PIEZO2 vs. IDR5^del^ p<0.0001. *n* number of cells per group is: GFP control 343, GFP 743, PIEZO2 733, IDR5^del^ 685. **c**, Cumulative distribution of the number of neurites per cell across GFP control, GFP, PIEZO2 and IDR5^del^ group. Total number of cells per group is: GFP control 836, GFP 854, PIEZO2 998, IDR5^del^ 810.

## Discussion

The original goal of this study was to examine the role of the seven intrinsically disordered intracellular domains of PIEZO2, which together account for almost one fourth of the channel. Our results highlight IDR5 as being particularly important for the function of PIEZO2. By demonstrating that IDR5 is required for force-from-filament gating of PIEZO2, but is dispensable for force-from-lipids activation of the channel, our study demonstrates that PIEZO2 can be activated by two different force transmission pathways. Moreover, by showing that the deletion of IDR5 renders PIEZO2 unable to inhibit neurite outgrowth, while it greatly though not completely reduces its sensitivity to mechanical indentation of the membrane and does not at all alter its sensitivity to membrane stretch, our data indicates that different types of physiological stimuli activate PIEZO2 via different force transmission pathways.

While our study eventually mainly focused on the role of IDR5, the initial characterization of the basal properties of the other IDR-deletions nevertheless provided interesting insights into their possible roles in PIEZO2 function. For example, we found that deletion of IDR3 significantly increases single channel conductance (Fig. 3c and d), which we believe was an indirect effect resulting from the dislocation of the plug domain (Fig. 3e), which – as we and others have previously shown – controls ion permeation through the lateral ion conducting portals of PIEZOs^25,26^. Since the plug domain and the lateral portals had, however, already been examined in great detail, we did not follow-up on the effect of IDR3 deletion in this study. Another observation we made was that deletion of IDR4 significantly increased stretch sensitivity of PIEZO2 (Fig. 2). This is very interesting, because a part of IDR4 is encoded by exon 18, which is missing in the PIEZO2 splice variants that are expressed in the lung and the bladder where PIEZO2 is mostly exposed to stretch rather than focal indentation of the cell membrane^51^. Hence, a role of IDR4 in fine-tuning PIEZO2 stretch-sensitivity would make perfect sense. Surprisingly, we did not observe any obvious functional deficits in PIEZO2 variants lacking IDR-1, −2, −6 or −7. This does of course not mean that these IDRs are completely dispensable since they might very well be important for functions that we did not examine here. Thus, they might serve as targets for modifying enzymes or as binding sites for auxiliary subunits that are only expressed in certain cell types but not in the N2a cells that were used in this study.

Two gating models – the force-from-lipids and the force-from-filament model – are commonly used to explain how mechanical forces exerted on a cell are transmitted to mechanically-gated ion channels embedded in the plasma membrane. The force-from-lipids model proposes that mechanical stimuli cause changes in the transbilayer pressure profile, which supposedly lead to conformational changes and thus activation of the channel, whereas the force-from-filament model suggests that mechanical forces are transmitted to the channel by the cytoskeleton either via direct interactions of via intracellular tethers^31,33^. While it was convincingly demonstrated that PIEZO1 is activated by force-from-lipids under experimental conditions in which the cytoskeleton is either completely missing (e.g. droplet bilayers, excised patches, membrane blebs) or partially detached from the membrane (cell-attached patches)^24,34,35^, it is completely unclear if additional mechanisms (e.g. force-from-filament) come into play in intact cells where the lipid bilayer and maybe even the channels themselves are tethered to the underlying actin cortex. Indeed, several other mechanically-gated ion channels, such as NOMPC, TMC1, ENaC, MEC4 and MEC10 were shown to require intra- and extracellular tethers for normal function^37,38,52,53^. TMC1 is particularly interesting, because despite the fact that it can be activated by force-from-lipids in liposomes, it still requires an intracellular tether for activation in intact cells^38,54^, which indicates that one cannot draw definitive conclusions about relevant in-vivo gating mechanisms based on inherent stretch sensitivity. There is also evidence for the requirement of an extracellular tether for normal mechanosensitivity of primary sensory afferents in which PIEZO2 mediates mechanotransduction^55,56^. Moreover, a contribution of the cytoskeleton to PIEZO channel activation seems highly likely, considering the observation that disruption of the cytoskeleton with Cytochalasin-D renders PIEZO1 and – as shown here – PIEZO2 less sensitive to mechanical cell indentation, but more sensitive to pressure-induced membrane stretch (Fig. 5a-g). For PIEZO1, the latter effect was previously shown to result from the removal of the mechanoprotective influence of the cytoskeleton, which renders the membrane more susceptible to pressure-induced stretch such that a given pressure pulse produces more stretch in Cytochalasin-D treated cells and thus larger currents. Our data shows that PIEZO2 is also ‘mechanoprotected’ by the cytoskeleton and can readily be activated by membrane stretch after Cytochalasin-D treatment, which indicates that PIEZO2 is also inherently sensitive to force-from-lipids (Fig. 2, 3, and 5).

Considering the aforementioned importance of intracellular tether for the function of other mechanosensitive ion channels, a straightforward explanation for the strong inhibitory effect of Cytochalasin-D treatment on poking-evoked PIEZO currents is that the majority of the channels that mediate whole-cell currents are activated by force-from-filament so that disruption of the cytoskeleton prevents force transmission to the channels, which would result in less activated channels and thus smaller current amplitudes. Most researchers in the field, however, favor an alternative explanation and proposed that the cytoskeleton indirectly transmits forces to the channel by generating local membrane tension via interactions with other membrane proteins, such that the ultimate stimulus that activates PIEZOs is force-from-lipids (Fig. 5l)^3,15,31–33^. Accordingly, this hypothesis implies that a channel with normal force-from-lipids sensitivity would also respond normal to membrane indentation in cells with an intact cytoskeleton. This is, however, not the case for IDR5^del^, which can readily be activated by pressure-induced membrane stretch (Fig. 2 and 3), which most likely depends on force-from-lipids gating, but only produces small currents in response to mechanical indentation of the cell (Fig. 1). Another implication of the latter hypothesis is that disruption of the cytoskeleton would reduce poking-evoked currents of channels that exhibit normal force-from-lipids sensitivity (see pathway 3 in Fig. 5l). This is also not the case for IDR5^del^, which appears to be completely insensitive to Cytochalasin-D-induced changes in cytoskeletal integrity (Fig. 5h-k). Thus, our data support the former gating model, which proposes that the great majority of the PIEZO2 channels that mediate poking-evoked whole-cell currents are activated by force-from-filament, and suggest that IDR5 mediates interactions between PIEZO2 and the cytoskeleton.

In summary our data shows that PIEZO2 is sensitive to both force-from-lipids as well as force-from-filament activation. Considering the differential effects of IDR5-deletion – i.e. removal of force-from-filament sensitivity – on stretch-evoked currents (no effect, Fig. 2 and 3), poking-evoked currents (partial reduction, Fig. 1) and neurite outgrowth (complete loss of inhibition, Fig. 6), it is tempting to speculate that different mechanical stimuli activate PIEZO2 via different force transmission pathways. Thus, we propose that pressure-induced membrane stretch in cell-attached recordings predominantly activates PIEZO2 via force-from-lipids, whereas mechanical indentation of the cell activates some channels via force-from-lipids and others via force-from-filament. Considering that changes in membrane tension, which are the direct consequence of membrane indentation, are locally restricted and are not propagated across the cell surface, we hypothesize that channels that are located in the close proximity of the stimulation probe and thus experience intense membrane stretch are predominantly activated by force-from-lipids, whereas channels that are located further away are predominantly activated by force-from-filament. Finally, considering that IDR5^del^ is normally sensitive to force-from-lipids but fails to significantly inhibit neurite outgrowth, we propose that cell-generated traction forces almost exclusively activate PIEZO2 via force-from-filament.

## Methods

### Cell culture and transfection

Neuro2A PIEZO1-Knockout cells (N2A-P1KO, gift from G.R Lewin) were grown at 37°C with 5 % CO2 in Dulbecco’s Modified Eagle Medium (DMEM) and optimal Minimal Essential Medium (opti-MEM) (1:1 mixture) with 10 % Fetal Bovine Serum (FBS), 2 mM L-glutamine and 1 % peniciline/streptomycine (all from Thermo Fisher). Cells were seeded on poly-L-lysine (Sigma) coated glass coverslips (for patch-clamp and immunocytochemistry), laminin coated coverslips (Neuvitro, for neurite imaging) or 35 mm 6 well plates (biotinylation assay). N2A cells were transfected one (neurite imaging, biotinylation) or two days (patch-clamp, immunohistochemistry) after plating using polyethylenimine (PEI, Linear PEI 25K, Polysciences). For one coverslip, 7 μl of a 360 μg/ml PEI solution is mixed with 9 μl PBS. Plasmid DNA is diluted in 20 μl PBS (0.6 μg/coverslip) and then added and mixed to the 16 μl PEI-PBS solution. After at least 5 minutes of incubation, 35 μl are added in one well and mixed by gentle swirling. For a 35 mm well, 1.5 μg DNA is used and PBS/PEI volumes are adjusted accordingly. 24 h later, the medium is replaced by fresh one. Cells are then used within 24 h (neurite imaging, biotinylation, patch-clamp) to 48h (patch-clamp, immunohistochemistry).

### Constructs and generation of PIEZO mutants

A PIEZO2-HA-IRES-GFP plasmid was previously created from mouse piezo2-pSPORT6 plasmid (gift from A. Patapoutian) and was used as the initial template to generate individual P2-IDR*X*^del^-HA-IRES-GFP construct, using a previously described strategy. Two AfeI restriction sites were sequentially added by two rounds of PCR-amplification on each side of individual IDR (see Fig. 1b and S1b for amino acid position) using KAPA HiFi polymerase (Roche). The PCR reaction was DpnI digested (New England Biolabs, 37°C, 1 h) and column purified (Macherey-Nagel) before being transform in electrocompetent Stbl4 bacteria (Invitrogen) and grown at 30°C for 48 h. After the two restriction sites were incorporated, plasmids were digested with AfeI (New England Biolabs) and gel purified to remove the cut IDR fragment. Then, the plasmid was re-ligated overnight at 16°C (ligase from Promega) and transform in Stbl4 bacteria. Selected clones were entirely sequenced to ensure that no other mutation was present.

### Neurite outgrowth assay

N2A-P1KO cells were prepared as described above. Cells were plated at a density of 5000 to 10000 cells per well containing one 12 mm laminin-coated coverslip and processed for transfection one day later. 24 h after transfection, cells were incubated with N2A medium without FBS and with Nerve Growth Factor (NGF, 100ng/ml, Sigma) to induce neurite outgrowth. Negative control coverslips were incubated instead with standard N2A medium. Neurites were imaged 30 to 40 h after using the GFP fluorescent signal from PIEZO transfected cells. An empty GFP vector was also used separately as a control. Fluorescent live images were acquired on an inverted microscope (IX70, Olympus) with a 20 x oil-immersion objective and visualized with an Imago-QE-Sensicam camera (PCO). Approximately 20 images were acquired per coverslips. One to three coverslips per constructs were used. The results presented here come from three independent experiments and transfection. Image analysis was manually performed on ImageJ, using the Region Of Interest (ROI) function to count and measure neurites. GFP-positive cell body perimeter was first marked, followed by the neurite and the side-branches.

### Whole-cell patch-clamp recordings

Mechanically activated currents were recorded at room temperature using EPC10 amplifier (HEKA) with Patchmaster and Fitmaster software (HEKA). The borosilicate patch pipettes (2–5 MΩ) were pulled with a Flaming-Brown puller (Sutter Instruments) and contained the following (in mM): 125 Kgluconate, 7 KCl, 1 MgCl_2_, 1 CaCl_2_, 4 EGTA, 10 HEPES, 2 GTP and 2 ATP (pH 7.3 with KOH). The control bath solution contained the following (in mM): 140 NaCl, 4 KCl, 1 MgCl_2_, 2 CaCl_2_, 4 glucose and 10 HEPES (pH 7.4 with NaOH). Cells were held at a holding potential of −60 mV and stimulated with a series of 13 mechanical stimuli in 0.4 μm increments with a fire-polished glass pipette (tip diameter 2-3μm) that was positioned at an angle of 45° to the surface of the dish and moved with a velocity of 1 μm/ms by a piezo driven micromanipulator (Nanomotor© MM3A, Kleindiek Nanotechnik). The evoked whole cell currents were recorded with a sampling frequency of 200 kHz and filtered with 2.9 kHz low-pass filter. Pipette and membrane capacitance were compensated using the auto function of Patchmaster. Leak currents before mechanical stimulations were subtracted off-line from the current traces. Recordings with excessive leak currents, unstable access resistance and cells which giga-seals did not withstand at least 7 consecutive mechanical steps stimulation were excluded from analyses.

The mechanical thresholds of the PIEZO2-mediated currents were determined by measuring the latency between the onset of the mechanical stimulus and the onset of the mechanically activated current. Current onset was defined as the point in which the current significantly differed from the baseline (< I_mean, baseline_ – 6 SD_baseline_). The membrane displacement at which the current was triggered was then calculated by multiplying the speed at which the mechanical probe moved (1 μm/ms) with the latency. The inactivation time constants (τ_inact_) were measured by fitting the mechanically activated currents with a single exponential function (C1+C2*exp(–(t– t0)/τ_inact_), where C1 and C2 are constants, t is time and τ_inact_ is the inactivation time constant. I/V curves and E_Rev_ were determined by changing the holding potential in −30mV steps (−60 to +60 mV) and by stimulating the cells with a fixed mechanical displacement that evoked a submaximal response. Only cells that withstand the full I/V protocol were used for E_Rev_ calculation.

To modulate PKA activity and assess its potential effect on PIEZO currents, N2A cells were first incubated the night prior the electrophysiological measurements with the PKA inhibitor KT5720 and the PKC inhibitor GF-109203 (both from Sigma, dissolved in DMSO and both used at a final concentration of 1 μM). Cells were further kept with GF and KT in the bath solution during the recordings. In order to stimulate PKA activity, cells that were previously exposed to GF and KT overnight were incubated 1 hour before the recordings with 1 μM GF, 300 μM of the PKA activator 8-Bromoadenosine 3’,5’-cyclic monophosphate (Sigma, dissolved in water) and without FBS. The drugs were also added to the standard electrophysiological bath solution during recordings. To investigate the contribution of cytoskeleton component in PIEZO currents, N2A cells were incubated with Cytochalasin-D for 15 minutes or with Nocodazole for 30min prior recordings (both from Sigma, dissolved in DMSO and both used at a final concentration of 1 μM). The drugs were also kept in the standard electrophysiological bath solution during the experiments.

### Single-channel recordings

Single-channel stretch-activated currents were recorded in the cell-attached configuration at room temperature using EPC10 amplifier (HEKA) with Patchmaster and Fitmaster software (HEKA). The borosilicate patch pipettes were coated with Sylgard (WPI) and fire polished (final resistance of 4–8 MΩ). The pipette solution contained the following (in mM): 130 NaCl, 5 KCl, 1 MgCl_2_, 1 CaCl_2_, 10 HEPES, 10 TEA-Cl (pH7.3 with NaOH). The bath solution contained (in mM): 140 KCl, 1 MgCl_2_, 2 CaCl_2_, 10 Glucose, 10 HEPES (pH7.4 with KOH). Pressure stimuli were applied with a 2ml syringe operated by a motorized device and measured with a custom-made pressure sensor or with the High-Speed Pressure Clamp (HSPC, ALA scientific). The evoked currents were recorded with a sampling frequency of 50 kHz and filtered with a 2.9 kHz low-pass filter. Pressure-response curves were evoked by a stepwise increase of negative pressure (3 seconds duration) with the cell being clamped at a holding potential of −100 mV. In response to repetitive and sustained pressure pulses, especially over −20 mmHg, PIEZO2 has the tendency to produce non-inactivating responses, making the determination of a potential “peak current” value difficult or impossible in most of the cases. Therefore, to accurately quantify stretch-activated PIEZO2 currents, we calculated the total charge transferred during the pressure stimulus (in pico Coulomb) through the determination of the area under the curve over the 3 seconds stimulus. Single-channel amplitudes at a given holding potential (−120 mV to −40 mV, 20 mV steps) were determined as the difference between the peaks of the gaussian fits of the trace histogram over a 500 ms segment. Unitary conductance was determined from the linear regression fits of the I/V plot of individual cells. Recordings with excessive leak currents (>4 pA) or unstable baseline were excluded from analyses. The effects of PKA and cytoskeletal modifying drugs were tested in the same conditions as for the whole-cell experiments.

### Immunocytochemistry

N2A cells were co-transfected with PIEZO2-HA-IRES-GFP or IDR^del^ constructs and with a plasmid encoding the red fluorescent protein mScarlet fused to a farnesylation signal sequence in its C-terminus to target it to the plasma membrane. Three days after transfection, cells were washed once with PBS and fixed with 4 % PFA for 10 minutes at room temperature, washed 3 times for 5 min with PBS and permeabilized for 1h at room temperature (permeabilization buffer: 2,5% donkey serum (Sigma), 1% BSA, 0.1% Triton X-100, 0.05% Tween-20, in PBS). Samples were then incubated overnight at 4°C with a 1:500 dilution of rabbit anti-HA antibody (Invitrogen) in PBS 1% BSA. After 3 washes of 5 min, cells were incubated for 1 hour with a 1:1000 dilution of AlexaFluor-647 donkey anti-rabbit (Life technologies, diluted in PBS 1% BSA) and washed 3 more times. Coverslips were mounted on slides with Fluoprobe mounting media that contain DAPI (Interchim). Confocal images were acquired with a SP8 confocal microscope (Leica) and a 63x oil-immersion objective. Images were analyzed offline with ImageJ.

### Membrane biotinylation assay

N2A cells were cultured and transfected as describe above. Two days after transfection, cells were washed twice with ice-cold PBS and treated for 30min with 0.6 mg/ml of biotin in PBS (EZLink Sulfo-NHS-LC-Biotin, Thermo Scientific). Next, an equivalent volume of glycin solution (100mM in PBS) was added to quench the remaining biotin. After 30 min, cells were washed twice with cold PBS and resuspended in 1 mL of PBS. To harmonize the number of cells between samples, turbidity at 600 nm was measured. Cells were then centrifuged for 2min at 4000rpm and lysed with RIPA buffer (50 mM Hepes, pH 7.4, 140 mM NaCl, 10% glycerol, 1% (v/v) Triton X-100, 1 mM EDTA, 2 mM EGTA, 0.5% deoxycholate, 10 mM PMSF and protease inhibitor mixture (Roche)) under gentle agitation for 1h at room temperature. Volume of RIPA buffer was by default 200μl for the sample having the highest optical density. The volume for the other samples was adjusted accordingly to obtain a similar cell concentration. Extracts were centrifuged at 11000 rpm for 15 min and the supernatant was kept. 10 μl of supernatant was taken and diluted with 10 μl of 4x Laemmli buffer (total extract fraction). Equilibrated Streptavidin resin (High Capacity Streptavidin Agarose, Thermo Fisher) was added to equal amounts of the remaining supernatants and incubated under gentle agitation overnight at 4 °C. After washing, biotinylated proteins were eluted by boiling the resin in 30 μl of 2 x Laemmli buffer for 5 min (biotinylated/membrane fraction), and subsequently analysed by SDS-PAGE and western blot.

### Western blotting

Biotinylated eluates and the corresponding total extract fraction were electrophoresed in 10 % polyacrylamide gels. Next, they were blotted onto a nitrocellulose membrane (0.4 μm, GE Health care) using a transfer buffer consisting of 30 mM Tris base, 190 mM glycine, and 20% methanol. After blocking at room temperature in TBS-T (20 mM Trizma base, 500 mM NaCl, 0.1% Tween-20) with 2% skimmed milk powder for at least 30min, membranes were incubated overnight with rabbit anti-HA (Sigma, dilution 1:1000 in TBS-T 5% skimmed milk) or rabbit anti-Beta tubulin (Sigma, used as a loading control, dilution 1:5000 in TBS-T 5% skimmed milk) at 4 °C. After washing with TBS-T, membranes were incubated with the secondary antibody for 1 hour at room temperature (Sigma, 1:10000, anti-rabbit HRP conjugated, diluted in TBS-T 5% skimmed milk). Finally, the immunoreactive bands were revealed using ECL Prime (GE Healthcare), visualized with the iBright1500 system and further analyzed with the iBright Analysis software (Invitrogen). The results presented here come from four independent experiments.

### Bioinformatics

To enable a better visualization of the location of the IDRs along the PIEZO2 blade, the unresolved IDRs were generated with SWISS-MODEL: they were first modeled using the full length PIEZO2 and the existing PIEZO2 cryo-EM structure (6KG7) as template. Then, the IDRS were removed from the model and added to the PIEZO2 structure (6KG7). All molecular images of PIEZO2 were generated with PyMOL 2.4.0 (Schrödinger, LLC). The mouse PIEZO2 amino acid sequence composition and properties were analyzed with Jalview 2.11.0. PKA site prediction analysis was conducted with NetPhos3.1^57^.

### Data analysis

Results were expressed as means ± s.e.m (unless otherwise noted). Statistical analyses were performed with Excel and Prism 8.0 (Graphpad). Data distribution was systematically evaluated and subsequent statistical tests were chosen accordingly. Two-tailed tests were used. Patch-clamp data were analyzed with FitMaster and Igorpro 8 (WaveMetrics). Statistical tests that were used, exact P-values and information about the number of replicates/cells are provided in each of the figure, in the corresponding legends or in the Source Data file. Symbols on graphics (* or #) indicate standard p-value range: *, p < 0.05; **, p < 0.01; ***, p < 0.001 and ns (not significant) p > 0.05.

## Data availability

Data supporting the findings of this manuscript and PIEZO mutant constructs are available from the corresponding authors upon request. A reporting summary for this article is available as a Supplementary Information file. The source data underlying figures and Supplementary Figures are provided as a Source Data file.

## Acknowledgements

This study was supported by the DFG grants LE3210/3-1 and SFB1158/1 to S.G.L. We thank Ms. Anke Niemann for technical assistance.

## Author contributions

C.V. cloned PIEZO2 mutants, performed and analysed patch-clamp recordings, biotinylation assays and neurite outgrowth assay and wrote the paper. I.S. cloned PIEZO2 mutants, performed and analysed patch-clamp recordings. J.M.J. and T.A.N. performed and analysed neurite outgrowth assays. W.N. performed immunocytochemistry. S.G.L. conceptualized the study, acquired funding, analysed the data, supervised the project and wrote the manuscript.

## Materials & Correspondence

Requests for materials and all other correspondence should be addressed to Stefan Lechner

