## Supplemental information for "“An intrinsically disordered intracellular domain of PIEZO2 is required for force-from-filament activation of the channel”"

**a**

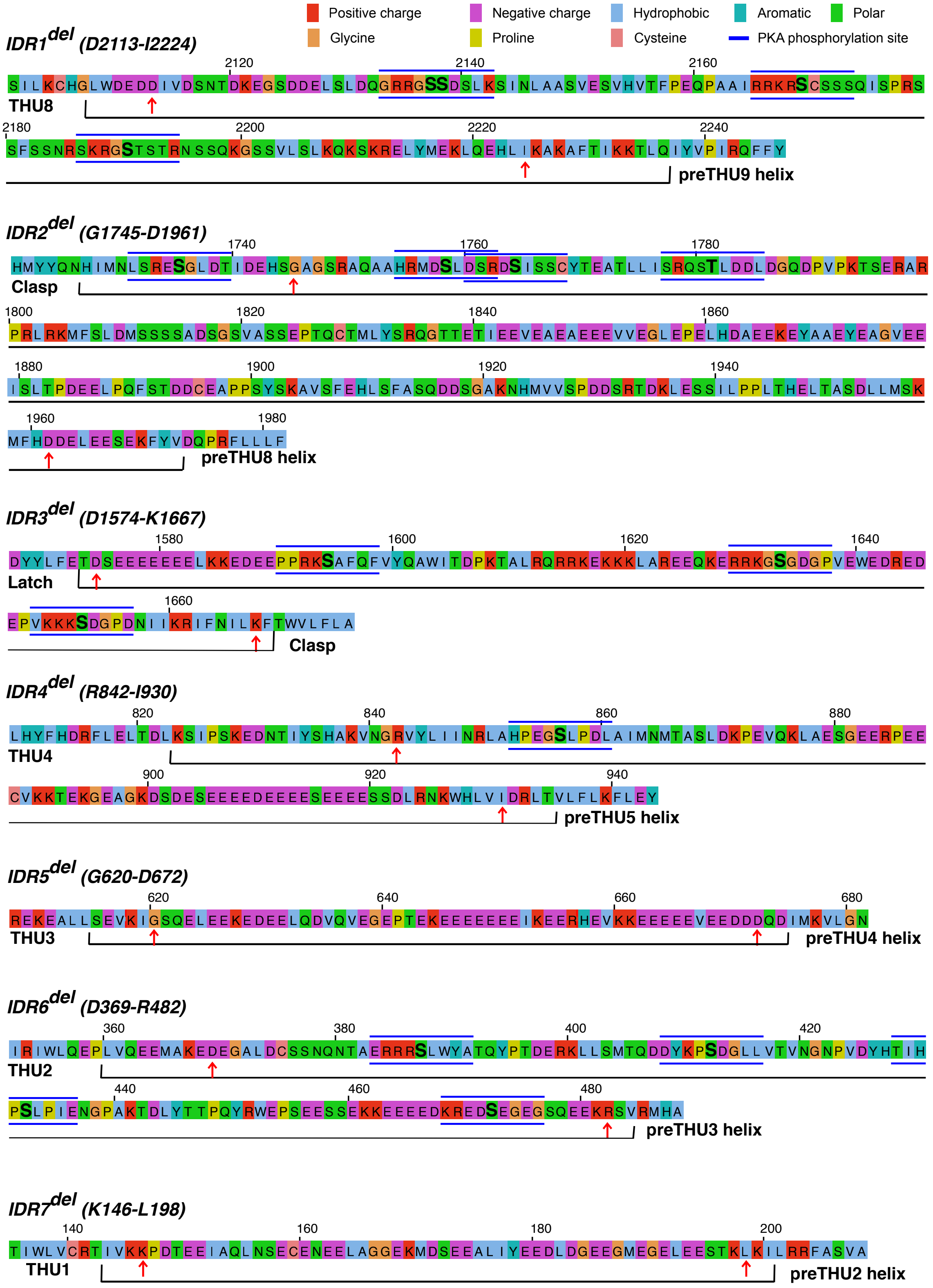


**Suppl. Fig. 1. Related to figure 1 and 4a**

**a,** Amino acid sequence details of the PIEZO2-IDR that were deleted in this study. Amino acid property is color-labelled based on Clustal code (indicated in legend), position and length of the intracellular IDR domain is indicated by the black bracket and the deleted region by the red arrows. PKA predictive sites are indicated by bold letter and blue lines.

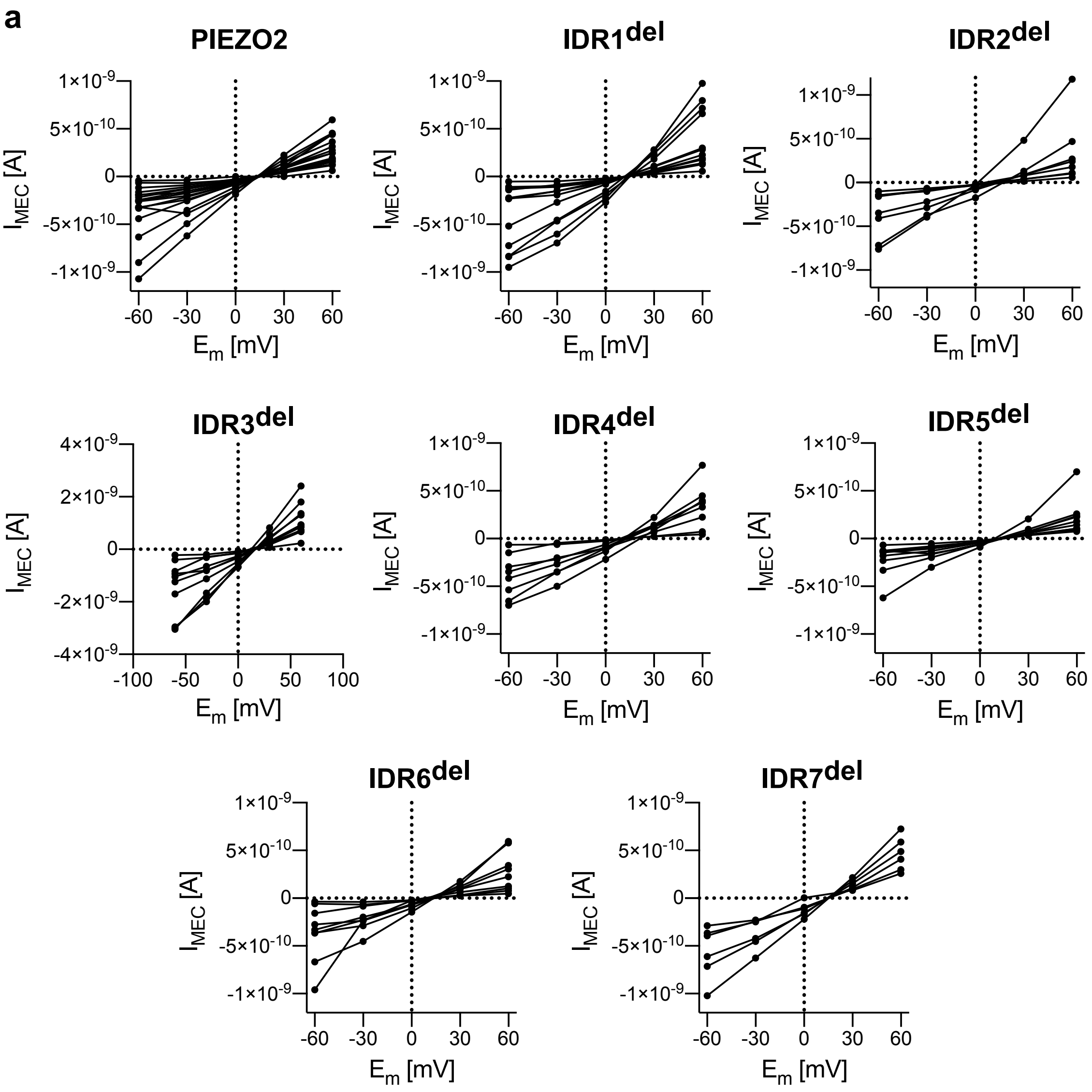


**Suppl. Fig. 2. Related to figure 1g**

**a,** Linear plots of the I-V relationships from individual cells in PIEZO2 and IDR$^{del}$ mutants. Symbols are individual peak current amplitude recorded at the indicated voltage. *n* number of cells per group are identical to fig. 1g.
